# *HtDCR-like1* regulates the development of structurally coloured cuticle by modulating cuticle chemistry and mechanical properties in *Hibiscus trionum*

**DOI:** 10.1101/2024.04.11.589056

**Authors:** Jordan Ferria, Siriel Saladin, Udhaya Ponraj, Raymond Wightman, Chiara Giorio, Chiara A. Airoldi, Beverley J. Glover

## Abstract

- Structural colours are a unique trait present in some animals, plants and even bacteria. In *Hibiscus trionum* flowers, they arise from nano-scaled cuticular ridges and may improve pollinator foraging efficiency. These ridges result from the buckling of the cuticle which is thought to be controlled by multiple parameters including anisotropic cell growth, chemical differentiation of the cuticle and formation of two layers of different mechanical properties.
- Here we investigate further the molecular and physical mechanisms by which structural colours are achieved in *Hibiscus trionum*. We produced two transgenic lines overexpressing *HtDCR-like1*, encoding a BAHD acyltransferase involved in cuticle synthesis. Transgenic lines showed impairment of the production of cuticle striations. Cuticle thickness, cuticle chemistry, cell elongation and cuticle stiffness were then investigated to identify the cause of failed buckling.
- We found that *HtDCR-like1* overexpression leads to modification of cuticle chemistry, including a decrease in detectable C_16_H_32_O_4_ (10,16-DHP) and a change to the measured Young’s moduli of the cuticle layers without alteration of the cell growth pattern.
- This work demonstrates the mechanisms by which gene regulation may control complex physical phenomena such as cuticle buckling via alteration of material properties of living tissues.

## Introduction

The control of physical parameters is crucial to the development of shapes and structures in living organisms. While the development of macroscopic structures can be explained at least partially by growth allometry, less is understood about the development of cellular and subcellular patterns. In plants these may consist of structures of various scales such as the micropapilae and nano-hairs of the sacred lotus leaf (*Nelumbo nucifera*) (Barthlott & Neinhuis, 1997; Zhang et al., 2016); the cuticular folds of *Paeonia mascula* (Moyroud et al., 2017)*, Hibiscus trionum* (Moyroud et al., 2017; Vignolini et al., 2015) and various other petal surfaces (Koch et al., 2013); the prismatic petal cells of *Eschscholzia californica* (Wilts et al., 2018) and *Gazania rigens* (which also displays cuticular folds) (Koch et al., 2013) and the conical petal cells of *Arabidopsis thaliana* as well as numerous other flowers (Gorton & Vogelmann, 1996; Irish, 2008; Whitney et al., 2009). Such patterns have been found in a wide variety of plants and perform a range of different ecological functions such as decreased or increased insect adhesion in *Hevea brasiliensis* (Surapaneni, et al., 2020; Whitney & Federle, 2013) and *Antirrhinum majus* (Whitney, Chittka, et al., 2009), respectively, and anisotropic wettability (Koch et al., 2013). In some cases, such as *Arabidopsis thaliana* striated conical cells, no ecological purpose is known (Panikashvili et al., 2009).

Some nano-scale surface patterns can produce optical effects that may act as signals. In *Hibiscus trionum*, observations of the proximal adaxial epidermis of the petals revealed the presence of a structurally coloured cuticle mainly characterised by the specific reflection of blue and UV-light in an angle dependant (iridescent) manner (Vignolini et al., 2015; Whitney et al., 2009). This optical effect was further shown to be visible to pollinators and sufficient to increase their foraging efficiency in a laboratory setting (Moyroud et al., 2017). The iridescent effect was shown to be the result of a diffraction grating-like structure formed from semi-ordered striations of the petal cuticle (Moyroud et al., 2017; Vignolini et al., 2015; Whitney et al., 2009). Although several studies have proposed potential physical mechanisms by which grating like surfaces are produced *in vitro* and *in vivo* (Airoldi et al., 2019; Airoldi et al., 2021; Bowden et al., 1998; Chen et al., 2021; Huang et al., 2017; Kourounioti et al., 2013; Lugo et al., 2023), the molecular regulation of the physical parameters leading to cuticular buckling remains unclear.

The cuticular patterns which function as diffraction gratings are continuous over multiple epidermal cells, which suggests that physical phenomena are involved. Experimental stretching of dissected tissue of non-striated immature *H. trionum* petals was sufficient to induce striations on the adaxial petal surface (Airoldi et al., 2021; Moyroud et al., 2022), which highlights the importance of tissue growth in cuticular buckling. Other studies have also shown the necessity of anisotropic cell elongation in the production of semi ordered cuticular micro pattern, e.g. on the petals of *Kalanchoe blossfeldiana* and *Eustoma grandiflorum* (Huang et al., 2017). However, the same mechanical perturbation was not sufficient to induce buckling of the abaxial petal cuticle of *H. trionum* (Airoldi et al., 2021), demonstrating that mechanical specialisation of the adaxial cuticle is necessary to allow buckling – a specialisation which would necessarily result from the activities of different genes in adaxial and abaxial epidermal tissues. Modification of the proximal adaxial cuticle chemistry resulting from *HtSHN3* over expression was sufficient to impair buckling without alteration of the anisotropic cell growth, indicating that cuticle chemistry is a crucial factor in cuticle buckling (Moyroud et al., 2022).

Studies of the *in vitro* mechanics of thin film buckling show that the presence of a bilayered system in which a thin and relatively stiff film tops a usually larger and more compliant substrate is necessary for the buckling of the upper layer (Bowden et al., 1998; Chen et al., 2021; Lugo et al., 2023). Several previous studies have demonstrated that the stiffness ratio between the substrate and the thin film plays a critical role in determining bilayer buckling and the wavelength of the resulting striations (Cao & Hutchinson, 2012; Cerda & Mahadevan, 2003). It has previously been suggested that a similar organisation of the cuticle could be critical for cuticle buckling in *Hibiscus trionum* (Airoldi et al., 2021, Lugo et al., 2023). In this petal diffraction grating system, the clear resolution of two separate layers is hampered by the relative continuity between the polysaccharides of the cell wall and the diversity of compounds associated with the cuticle. Transmission electron microscopy of cross sections of the adaxial petal part distinguished at least two regions based on their electron density: a thin outer layer of about 200 nm<u>, usually referred to as the cuticle film</u>, and a thicker inner region, usually referred to as the extracellular matrix or <u>cuticular layer</u> (Vignolini et al., 2015, <u>Lugo et al., 2023</u>). Fat-red staining also supported the lipidic nature of the outer 200 nm layer (Moyroud et al., 2022). The outer layer is interpreted as the cuticle proper and is predominantly composed of cuticular material, while the inner layer is likely composed of both cuticular and cell wall compounds. The molecular mechanisms allowing the development of a stiffer cuticle film remain unclear. However, specific analysis of the chemistry of the cuticle film with Liquid extraction surface analysis coupled with high precision mass spectrometry (LESA-MS) shows that the capacity of the cuticle to buckle is tightly linked to its chemistry, supporting the idea that cuticle chemistry determines the stiffness of the cuticle and that cuticle chemistry is tightly regulated to allow cuticle buckling (Giorio et al., 2015; Moyroud et al., 2022).

Aside from residual cellulose polymers, the cutin network constitutes one of the major macromolecules of the cuticle and consists of polymerised aliphatic acids, often of 16 and 18 carbon residues, such as 10,16-dihydroxypalmitate (10,16-DHP), mostly linked together by ester bonds, as well as other linkers such as glycerol and triacylglycerol (Fernández et al., 2016). Chemical alteration of the cuticle in *Hibiscus trionum* prevented buckling properties of the cuticle upon artificial stretching, showing direct alteration of the mechanical properties of the cuticle (Moyroud et al., 2022). The mechanical properties of tomato fruit cuticle were also found altered by the modification of the cuticle chemistry (Isaacson et al., 2009). It could therefore be hypothesised that the modulation of the chemistry of the cuticle film in *Hibiscus trionum* results in an alteration of the cutin network, which in turn will alter its Young’s modulus and therefore its buckling capacity.

The production of this cutin network is the result of the expression and coordination of numerous transcription factors and downstream structural proteins such as SHN1/2/3 (AP2/EREBP transcription factors) (Aharoni et al., 2004; Shi et al., 2013, 2011), DCR (a BAHD acyltransferase) (Mazurek et al., 2017; Panikashvili et al., 2009; Rani et al., 2010), GPAT6 (a glycerol-3-phosphate acyltransferase) (Li-Beisson et al., 2009; Petit et al., 2016), CYP77a6 (involved in mid-chain hydroxylation of hydroxypalmitic acid) (Lashbrooke et al., 2015), ABCG11 and LTPs (involved in the transfer of cutin monomers towards the extracellular environment) (Bird et al., 2007; Jacq et al., 2017), and CD1 (catalysing cutin polymerisation) (Fich et al., 2016; Yeats et al., 2012).

Although the DCR (DEFECTIVE IN CUTICULAR RIDGES) protein has been identified as an acyltransferase (Molina & Kosma, 2015; Panikashvili et al., 2009; Rani et al., 2010), its biological role remains unclear. It was first suggested that diacylglycerol was the main substrate of DCR (Rani et al., 2010), but later reports suggested that palmitic acid or hydroxypalmitic acid could also act as substrates of DCR (Molina & Kosma, 2015). In the latter scenario, hydroxypalmitic acid acylation was suggested as potentially necessary for the subsequent mid-chain hydroxylation by CYP77a6 (Molina & Kosma, 2015), yielding 10,16-dihydroxypalmitate. It has also been suggested that DCR intervention is necessary for the production of triacylglycerols that are subsequently integrated into the cutin network (Mazurek et al., 2017; Rani et al., 2010). Despite the lack of consensus, DCR appears to play a central role in the synthesis of cutin and its monomers, potentially via the production and maturation of dihydroxypalmitic acid and the associated triglycerides. Furthermore, it has been demonstrated in *Arabidopsis thaliana* petals that DCR is important for the integration of 10,16-DHP into the cuticle and that it also mediates the structure of the cuticle regardless of its composition (Mazurek et al., 2017). The loss of function *dcr1* mutant of *A. thaliana* is associated with loss of striations on the cuticle of the abaxial and adaxial cells of the petals and a depletion in 10,16-dihydroxypalmitate (10,16-DHP), which is consistent with a potential role in its synthesis (Panikashvili et al., 2009). LESA-MS (liquid surface extraction analysis coupled to high-resolution mass spectrometry) on structurally coloured and non-structurally coloured species of *Hibiscus* and varieties and transgenic lines of *Hibiscus trionum* also revealed a strong association between 10,16-dihydroxypalmitate and the presence of cuticular striations (Moyroud et al., 2022). Therefore, alteration of the cuticle chemistry and of the cutin network through the activity of DCR is predicted to result in a failure of cuticular buckling.

Other genes have also been shown to play a role in cuticle buckling. Alteration of the synthesis of the cutin polymer by overexpressing the cutin polymerase *HtCD1* was correlated with partial impairment of *H. trionum* cuticular striations, supporting the potential role of the cutin network in the production of striations (similar results were obtained with *HtDEWAX* over expression transgenic lines) (Moyroud et al., 2022). Together, these results suggest potential molecular routes by which the physical parameters of the cuticle can be adjusted to induce its buckling. However, although numerous clues suggest that cuticle chemistry might directly regulate its mechanical properties, the link between cuticle chemistry, its Young’s modulus and the production of structural colours has yet to be demonstrated directly.

In this paper we investigate the partial loss of structurally coloured cuticle in two *Hibiscus trionum* transgenic lines overexpressing *HtDCR-like1*, a putative orthologue of *AtDCR.* We compare the chemical and physical parameters of striated and non-striated cuticles in both WT and transgenic lines, in an effort to bridge the molecular regulation of cuticle synthesis, its chemistry and the resulting physical parameters, which, in turn, determine whether the cuticle will buckle and produce structural colour.

## Material and methods

### Plant material

Wild-type (WT) plants of *Hibiscus trionum* were from the Glover lab stock line obtained from Cambridge University Botanic Garden. Following transformation, a total of five transgenic plants from two independent transgenic lines were obtained: three sister plants from transgenic line 1 (*35S:HtDCR-like1* Independent Line 1): P1, P2 and P3, and two sister plants from transgenic line 2 (*35S:HtDCR-like1* Independent Line 2): P4 and P5.

### Cloning and Plant transformation

The *HtDCR*-*like1* coding sequence was isolated from *Hibiscus trionum* proximal petal region cDNA. cDNA was obtained following RNA extraction using the Sigma Aldrich® Spectrum™ plant total RNA kit and subsequent Thermo Fischer Scientific® DNase I treatment at 37 °C for 30 min., and reverse transcription using oligodT primers and BioScript™ reverse transcriptase with incubation at 42 °C for 45 min.

Isolation of *HtDCR-like1* was performed with tailed primers (see supplementary table S1) designed against a *de novo* assembly of the *Hibiscus trionum* petal transcriptome (Ferria, 2022) and containing SapI, partial BsaI and SmaI or EcoRI restriction sites necessary for cloning into pUAP4 as described in (Frangedakis & Sauret-Gueto, 2020) and subsequent cloning into our expression vector.

Overexpression vector pEM110 was built from a pGREENII backbone (Hellens et al., 2000) and contains a double CaMV 35S promoter (Moyroud et al., 2022). The resulting vector was then used to perform an *Agrobacterium tumefaciens* mediated transformation of *Hibiscus trionum* as detailed in (Moyroud et al., 2022).

### Protein and nucleic acid alignment

The *Arabidopsis thaliana DCR* gene (also known as *AtPEL3*) sequence is referenced in (Panikashvili et al., 2009) and its protein and transcript sequences were retrieved from the TAIR database under the accession number *At5g23940*. Protein alignment and visualisation was performed using the online T-coffee v11 platform (https://tcoffee.crg.eu) using the PSI-coffee homology extension and Clustal omega algorithm with default parameters.

Protein alignment for phylogeny analysis was performed using Mview (v 1.67) with default parameters and the Clustal omega algorithm. Subsequently, PartitionFinder (v 2.1.1) was used without manual partitioning implementation for optimal sequence alignment, and RAxML-ng (v 1.1.0) with the JTT and DCMUT+G models was used for the resolution of the DCR phylogeny. Finally, FigTree (version 1.4.4) was used for phylogeny visualisation.

Transcript alignment was performed using MACSE and RAxML-ng using the GTR+G model for phylogeny resolution and FigTree was used to visualise the phylogenetic tree.

### Chemical analyses

Chemistry of the adaxial cuticle was analysed using liquid extraction surface analysis coupled to high-resolution mass spectrometry (LESA-MS) as described in (Giorio et al., 2015). In total, six flowers were analysed: three from each transgenic line. WT chemical profiles were acquired following the same protocol and were published in (Moyroud et al., 2022) for a total of four flowers.

A non-polar (chloroform:acetonitrile:0.1% formic acid water (49:49:2)) and a polar (acetonitrile:0.1% formic acid water (9:1)) solvent were used on the cuticle of the adaxial petal surface of both proximal and distal regions of each sample, except on the proximal petal parts of samples WTb and WTd, which were only analysed with non-polar solvent. Additional details on the mass spectrometry analysis and the data processing can be found in (Giorio et al., 2015; Zielinski et al., 2018).

Mass spectrometry allowed for the detection of 110 compounds over all six transgenic and WT samples. Multiple measurements were performed per flower yielding a percentage of detection of each compound for each flower giving a chemical profile specific to each sample. The R package ‘ade4’ was used for Principal Component Analyses (PCA). Statistical comparisons between transgenics and WT was performed using a non-parametric pairwise Wilcoxon test with a Benjamini and Hochberg’s correction for the C_16_-C_18_ cuticle compounds.

### Imaging

CryoSEM and cryofractures were carried out on a Zeiss EVO HD15 equipped with a Quorum PPT010 cryo preparation system according to the protocol described in (Wightman et al., 2017) with the following modifications: (i) petal surfaces were coated with 5 nm gold-palladium, (ii) cryofractures were coated with 3 nm of gold-palladium, (iii) all imaging used 6 kV gun voltage and an SE2 detector. Samples were prepared at the microscopy facility of the Sainsbury Laboratory Cambridge University (SLCU).

Optical microscopy was conducted on a Keyence microscope equipped with either a 20x-200x or a 100x-1000x objective. Prior to observations, petals were stuck flat onto a petri dish using double sided tape.

Macroscopic pictures of *Hibiscus trionum* flowers were taken using a Samsung Galaxy S8 cell phone: dual Pixel 12 Megapixels autofocus, OIS (Optical Image Stabilization), F1.7 aperture with a pixel size of 1.4 µm and a sensor size: 1/2.55.

### Cuticle thickness and aspect ratio

Thickness of the cuticle film and aspect ratio of the striations were determined on SEM pictures of cross sections of the proximal adaxial petal surface using ImageJ. Aspect ratio (AR) and the thickness of the cuticle film were determined together for each cell that was analysed. The mean aspect ratio of the striations of a cell was determined by dividing the average height (h) by the average wavelength (λ) measured (AR= h/λ). An AR of ‘0’ was automatically inferred for smooth cells.

AR and thickness of the cuticle film were assessed in 52 WT cells over 2 different individuals (30 + 22 cells), 82 cells of 3 individuals of the *35S:HtDCR* IL 1 (32+31+19) transgenic line and 73 cells of the 2 individuals of the *35S:HtDCR* IL2 (26+47) transgenic line. Image resolution and sample preparation did not allow for a reliable measure of the cuticle film for 6 of these cells, reducing the sample size to 52, 77 and 72 cells per genotype, respectively.

### Cell size measurement

Cell size was measured on optical microscopy pictures taken with the Keyence microscope at a magnification of 300x at various developmental stages: before (10 mm buds), during (12.5-13 mm buds) and after (20 mm and open flowers) the development of striations. For 20 mm buds and open flowers of the transgenic lines, only smooth or abnormally striated cells were measured over a transect going from one side to the other of the petal for the transgenic lines. For 10 mm buds, 120 cells were measured over 4 randomly selected petals for each bud; for 12.5-13 mm buds, 180 cells were measured over 3 petals for each bud; for 20 mm buds, 120 cells were measured over 3 petals for each bud and finally for open flowers, 120 cells were measured of 3 petals per flower. For WT plants two buds/open flowers from two biological replicates were measured. For *35S:HtDCR-like1* transgenic line 1, one bud/open flower was measured for each of the three plants cultivated. For *35S:HtDCR-like1* transgenic line 2, one bud/open flower was measured from each of the two plants cultivated for each developmental stage, except for the 20 mm bud which was only measured in one of the two plants cultivated.

### Stiffness measurements

Bud stage 12.5–13mm was used from WT, *35S:HtDCR-like1* transgenic line 1 and *35S:HtDCR-like1* transgenic line 2 to calculate the stiffnesses of the cuticle film and the underlying cuticular layer using Atomic Force Microscopy (AFM). From each line three flower buds were used and from each flower bud three samples were measured at three different random positions both before and after mechanical removal of the cuticle film. Samples were prepared and placed on 3 % agarose and measured in water with 0.005 M sorbitol as described by (Airoldi et al., 2024). Indentation depth was kept below 29 nm, which is below 1/10 of the thickness of both the cuticular film and cuticle layer.

Force displacement curves were measured using AFM JPK Nano Wizard with a cantilever of tetrahedral shape (ATEC-CONT Au10 S/N: 74454F6L1180) as described in (Airoldi et al., 2024). For each new cantilever the spring constant was calibrated using the thermal noise method in the air. We used five cantilevers in our experiment with spring constant of 0.26 N/m, 0.25 N/m, 0.26 N/m, 0.24 N/m and 0.20 N/m. The sensitivity was calculated in water at the beginning of every set of measurements. The measured values for each experiment ranged between 50 and 80 nm/V. JPK software (JPK 00806) was used in force mapping mode to record force displacement curves with settings as described in(Airoldi et al., 2024). The collected measurements were analysed using JPK SPM Data Processing BRUKER software and Young’s moduli values were calculated as (Airoldi et al., 2024). Examples of Young’s modulus frequency curves and their associated AFM maps are provided for each genotype before and after cuticle film removal in supplementary figure 1 (S1)

### qPCRh

RNA was extracted from the proximal region of open flowers of a WT individual and five transgenic plants using the Sigma Aldrich® Spectrum™ plant total RNA kit with 15 min. on column DNase treatment. Reverse transcription was performed using the BioScript™ reverse transcriptase for 45 min. at 42 °C using oligodT primers. Quantitative PCR was then performed using the Luna® Universal qPCR Master Mix for a final volume of 10 μL. ActinS4 was targeted as a housekeeping gene using primers as in (Moyroud et al., 2022). Primer efficiency of the q-PCR *HtDCR-like1* primers (see supplementary table S1) was 104.8%. Relative *HtDCR-like1* expression for each plant was presented as the 2^−ΔCt^_transgenic_−2^−ΔCt^_WT_ score. Standard error of mean was calculated as follows: *SEM* = *Ln*(2) ∗ *E* ∗ 2^−ΔCt^, with *E* = 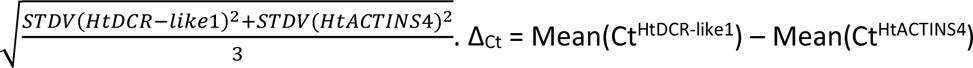

## Results

### The HtDCR-like1 protein contains signature motifs of the HXXXD/BAHD acyltransferase protein family

A BLAST search for *AtDCR* in a *Hibiscus trionum* petal transcriptome (Ferria, 2022) and a full genome assembly (kindly made available by Edwige Moyroud, Sainsbury Laboratory Cambridge University) yielded two *HtDCR* loci named *HtDCR-like1* and *HtDCR-like2*. *HtDCR-like1* and *2* genes were subsequently blasted using NCBI BLAST against the genomes of *Cucumis sativa* (Cucurbitaceae), *Theobroma cacao* (Malvaceae), *Gossyium raimondii* (Malvaceae), *Arabidopsis thaliana* (Brassicaceae) and *Brassica oleracea* (Brassicaceae), yielding only one locus from each genome. We generated hypotheses about the putative function of *Hibiscus trionum* DCR proteins by performing multiple sequence alignment between AtDCR, HtDCR-like1, HtDCR-like2, *Gossypium raimondii* DCR-like, *Theobroma cacao* DCR, *Brassica oleraceae* DCR and *Cucumis sativa* (out group) DCR protein sequences (Fig. **1a**). Analysis of the protein sequence revealed the presence of two highly conserved motifs: HXXXD and DFGWD, characteristic of the HXXXD/BAHD acyltransferase protein family, in all seven sequences. Furthermore, HtDCR-like1 and AtDCR/PEL3 were found to be over 69% identical, suggesting a similar function between HtDCR-like1 and AtDCR proteins (see supplementary figure S1a). A phylogenetic analysis based on the same protein sequences (Fig. **1b**) and their equivalent coding sequences (see supplementary figure S2) was also conducted. The observed phylogenetic tree matches the hypothesis of a single DCR orthologue in the Malvaceae genomes, with a recent duplication in the lineage including *H. trionum.* The same topology was produced by both transcript-based and protein-based phylogenies, which is consistent with the possibility of protein functional conservation during evolution. This similarity between AtDCR and HtDCR-like proteins suggests that HtDCR-like proteins may be involved in the production of structurally coloured cuticle in *Hibiscus trionum,* similar to the role of *AtDCR* in the development of cuticular striations on the *Arabidopsis thaliana* adaxial sepal epidermis (Panikashvili et al., 2009). Since *HtDCR-like1* and *HtDCR-like2* are 92.94% identical (see supplementary figure S2a), we suspected that there would be redundancy in their regulation of cuticle chemistry and that any overexpression phenotype would be very similar. We thus only transformed *Hibiscus trionum* with a *35S:HtDCR-like1* construct.

**Figure 1:**
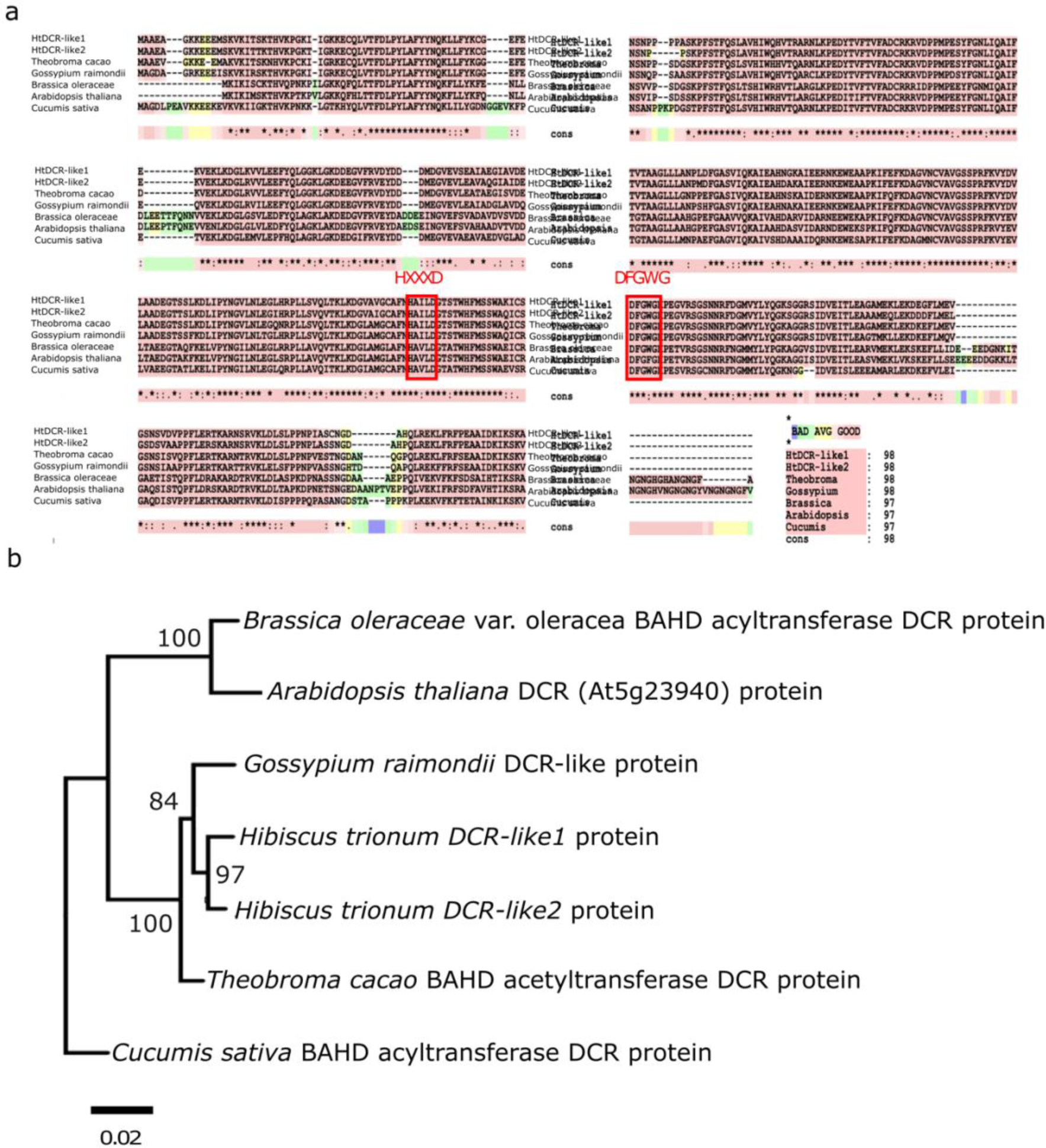
Protein sequence alignment suggests a similar function between HtDCR and AtDCR. **a**) Alignment of the DCR and DCR-like proteins performed with T-coffee v.11 using Clustal omega algorithm and default parameters. Green and blue highlightings show a relatively ‘bad’ alignment, yellow an ‘average’ alignment and red a ‘good’ alignment. Overall alignment score is 984. Red frames highlight the two signature domains of the BAHD acyltransferase protein family: HXXXD and DFGWG (Molina et al., 2015). **b**) Bootstrapped phylogeny based on DCR and DCR-like amino-acid sequences using the JTT model of raxml (similar results were obtained using the DCMUT+G model, data not shown).

### Overexpression of *HtDCR-like1* leads to partial loss of cuticular striations in *Hibiscus trionum*

To investigate the role of *HtDCR-like1* in the production of structural colours, we overexpressed *HtDCR-like1* in *Hibiscus trionum.* Two independent transgenic lines (*35S:HtDCR-like1* IL1 and *35S:HtDCR-like1* IL2) were generated and in total five plants were analysed (three sister plants originating from the transgenic line 1 (IL1) (P1, P2 and P3) and two sister plants originating from transgenic line 2 (IL2) (P4 and P5). Quantitative-RT PCR results indicate a significant increase in *HtDCR-like1* transcript level in the proximal petal part of open flowers in all the transgenic plants compared to that of WT plants (Fig. **2a**).

**Figure 2:**
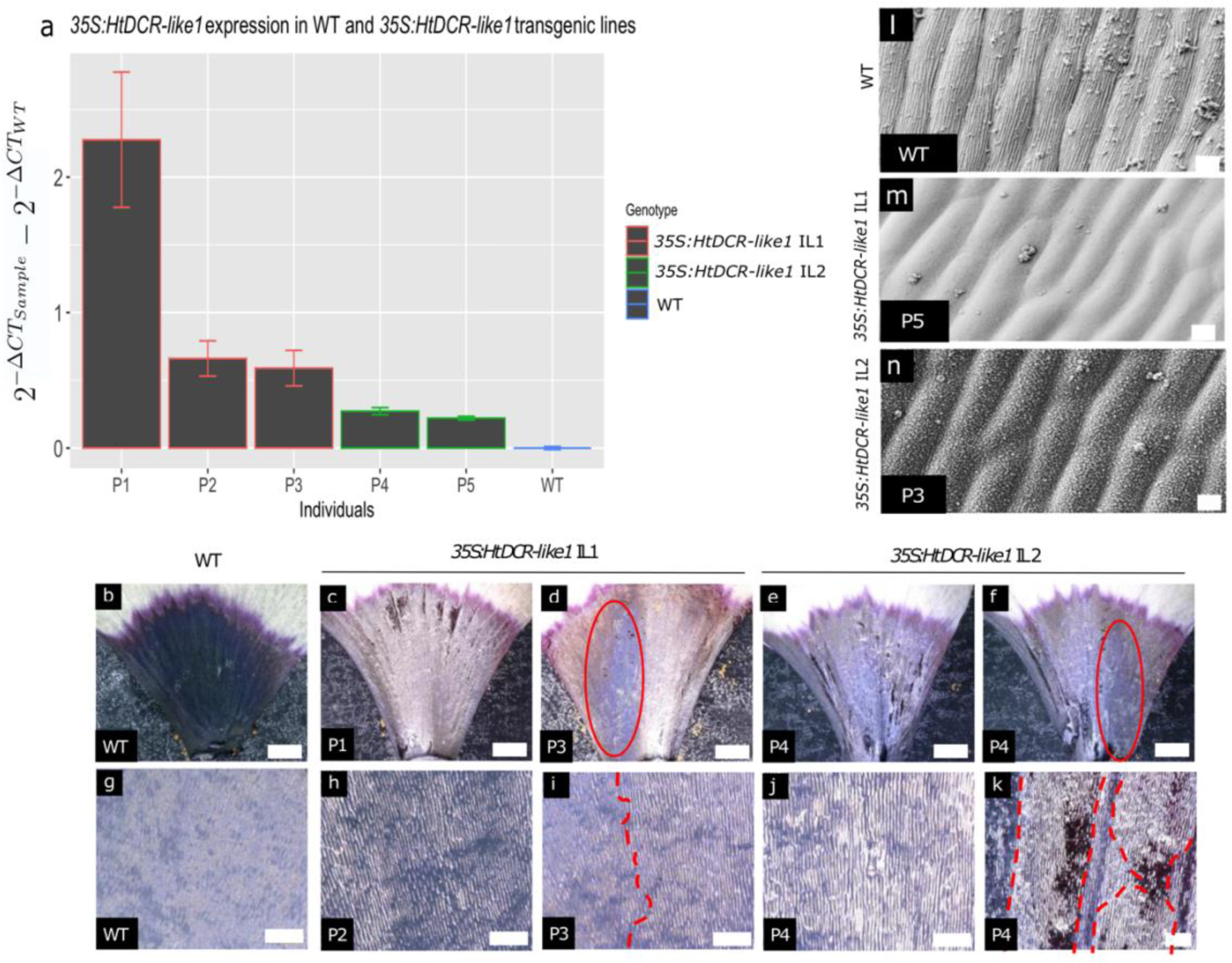
Overexpression of *HtDCR-like1* results in partial alteration of the petal cuticular striations. **a**) Relative expression of *HtDCR-like1* measured in the proximal petal region of fully open *Hibiscus trionum* presented as 2^−Δ*Ct*^_*transgenic*_ − 2^−Δ*Ct*^_*wt*_. Δ^Ct^ being the difference of Ct observed between *HtDCR-like1* and *HtActinS4* (housekeeping gene). **b-f**) Optical microscopy of the proximal adaxial petal surface (scale bar = 1 mm). **g-k**) Optical microscopy pictures of the centre of the adaxial proximal petal surface (scale bar = 200 μm). Circles showing striated and blue-reflecting cuticle. Dashed lines delimiting striated and non-striated regions. **l-n**) SEM of the adaxial proximal petal surface (scale bar = 10 μm).**c**, **d**, **h**, **i** and **m**) *35S:HtDCR-like1* IL1. **e**, **f**, **j**, **k** and **n**) *35S:HtDCR-like1* IL2. **b**, **g** and **l**) WT. **c**) Plant P1. **h**) Plant P2. **d**, **i** and **m**) Plant P3. **e**, **f**, **g** and **k**) Plant P4. **n**) Plant P5.

Phenotypic analyses were performed on the proximal petal region in both WT and *HtDCR-like1* overexpression transgenic lines using optical and electron microscopy (Fig. 2 and supplementary figure S3). Comparison of transgenic and WT flowers revealed an alteration of the blue hues in the proximal part of the petal (supplementary figure S3), which is consistent with complete or partial loss of cuticular striations. Optical microscopy confirmed the presence of patches of smooth cuticle and even a complete loss of striations in some petals (Fig. **2b-k**). Electron microscopy confirmed the presence of smooth cuticle in various amounts (Fig. **2l-n**, supplementary figure S3).

### *HtDCR-like1* overexpression affects striation aspect ratio without alteration of the thickness of the cuticle film

The optical microscopy observations of the striated regions of *HtDCR-like1* over expression lines suggested that some cells displayed intermediate phenotypes with a lack of consistency in the aspect ratio of their striations. This was confirmed by electron microscopy performed on fractures of the adaxial petal surface (Fig. 3 **a-h**). The aspect ratio of the striations in both WT and overexpression lines was quantified on cyroSEM fracture images of cross sections of the cells of the adaxial epidermis of the petal (**Fig. 3i** and **j**).

**Figure 3:**
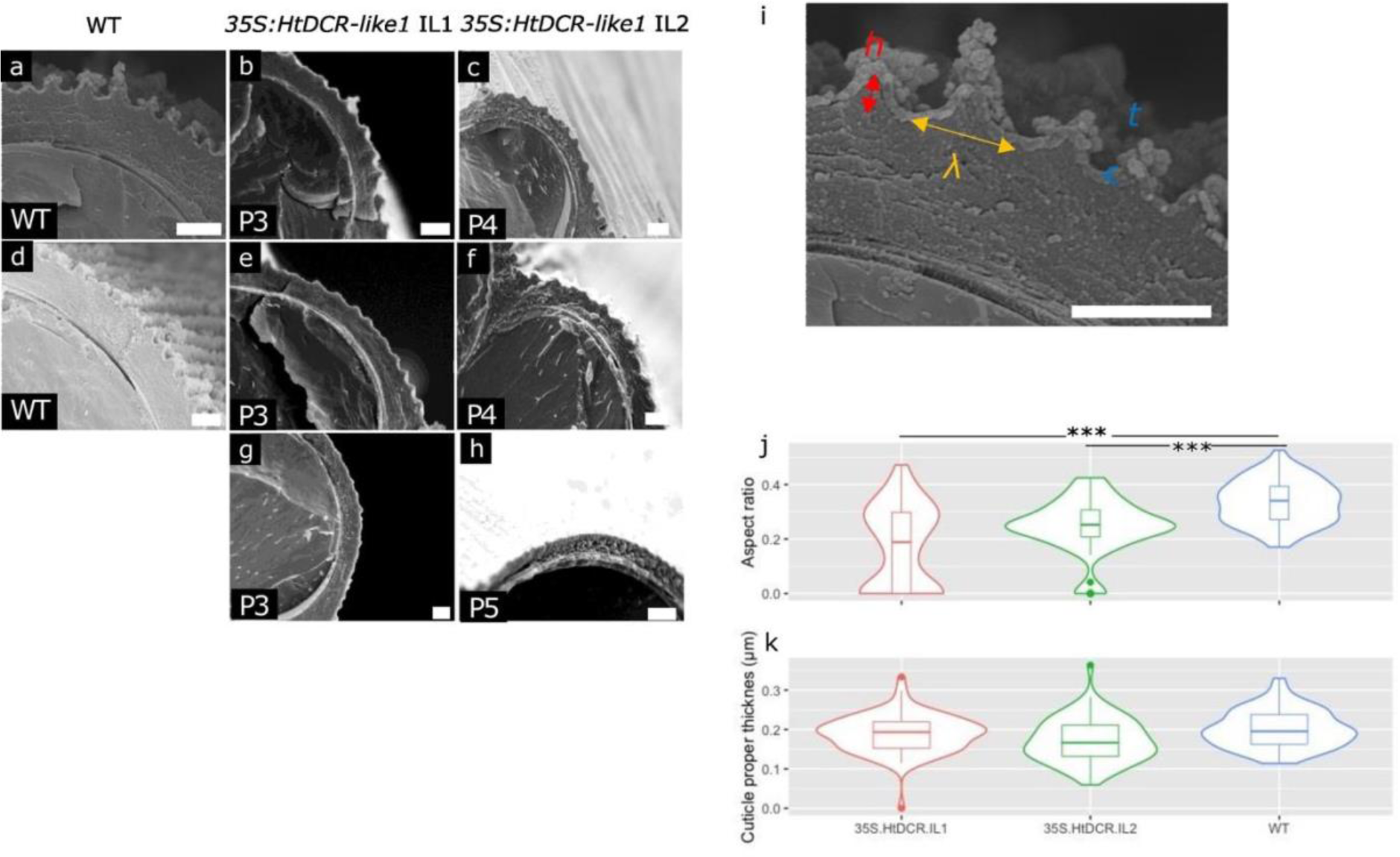
Overexpression of *HtDCR-like1* results in alteration of the aspect ratio of the cuticular striations without alteration of the thickness of the cuticular film. **a-f**) Scanning electron microscopy of the adaxial proximal petal cuticle observed on cryofractured samples. Scale bar = 2 μm. **a** and **d**) WT *Hibiscus trionum*. **b**, **e** and **g**) Plants of the *35S:HtDCR-lik*e*1* IL1 transgenic line. **c**, **f** and **h**) Plants of the *35S:HtDCR-like1* IL2 transgenic line. **i**) parameters measured on cross sections of *Hibiscus trionum* adaxial petal epidermis: h = striation height; λ = striation wavelength; t = thickness of the cuticular film. **j**) Average aspect ratio (h/λ) measured for each cell on SEM pictures of cross sections of the proximal adaxial petal cuticle. Kruskal Wallis test and pairewise Wilcoxon test with Benjamini Hochberg’s correction showed statistical differences: *** ⇐⇒ 0.001 > p-value. **k**) Cuticular film thickness measured per cell on SEM pictures of cross sections of the proximal adaxial petal cuticle. Anova test showed no significant difference.

In both sets of transgenic lines, overexpression of *HtDCR-like1* results in a significant reduction of the aspect ratio of the remaining striated cells (Fig. **3j**).

Alteration of the aspect ratio of striations and a loss of striations may be the result of an alteration of the physical parameters required for thin film buckling. In theory, the thickness of the cuticle film can affect the aspect ratio of the striations and potentially induce a failure of buckling (Biot, 1957; Genzer & Groenewold, 2006; Lugo et al., 2023). Thus, the thickness of the cuticle film in the same cells was also assessed (Fig. **3k**). No significant difference was recorded between the thickness of the cuticle film of the WT and that of the two transgenic lines, despite significant variation in the aspect ratio of the striations, suggesting that the observed phenotype is not the result of alteration of the thickness of the cuticle film.

### *HtDCR-like1* overexpression did not induce significant alteration of the cell growth pattern

Alteration of the growth pattern of cells has also been suggested to have an impact on the development of striations (Huang et al., 2017; Kourounioti et al., 2013). To investigate cell growth, we measured cell size at 4 different stages (**Fig. 4a** and **b**). Although relevant differences in cell size were measured between WT and transgenic buds and flowers, no clear alteration of the overall growth pattern could be deduced from cell size only. To compare the actual cell growth between pre-striation and post-striation stages, we calculated the average cell deformation in length and width (λ_1_ and λ_2,_ respectively), accounting for the two-dimensional strains, at the beginning of development: between 10 mm buds and 12.5-13-mm buds, and for overall development between 10 mm buds and fully mature flowers (Fig. **4c**). Most samples are contained within WT λ_1_ and λ_2_ values during both early and overall growth and display transition between the strains measured at the early stages of development and those measured on the overall petal growth which are similar to those of WT1. The evolution of λ_1_ measured in both transgenic lines between the early developmental stages of the bud and on the overall flower development is consistently larger than that of WT2, suggesting that the cell elongation along the longitudinal axis of the petal in the transgenic lines should be sufficient to induce cuticle buckling.

**Figure 4:**
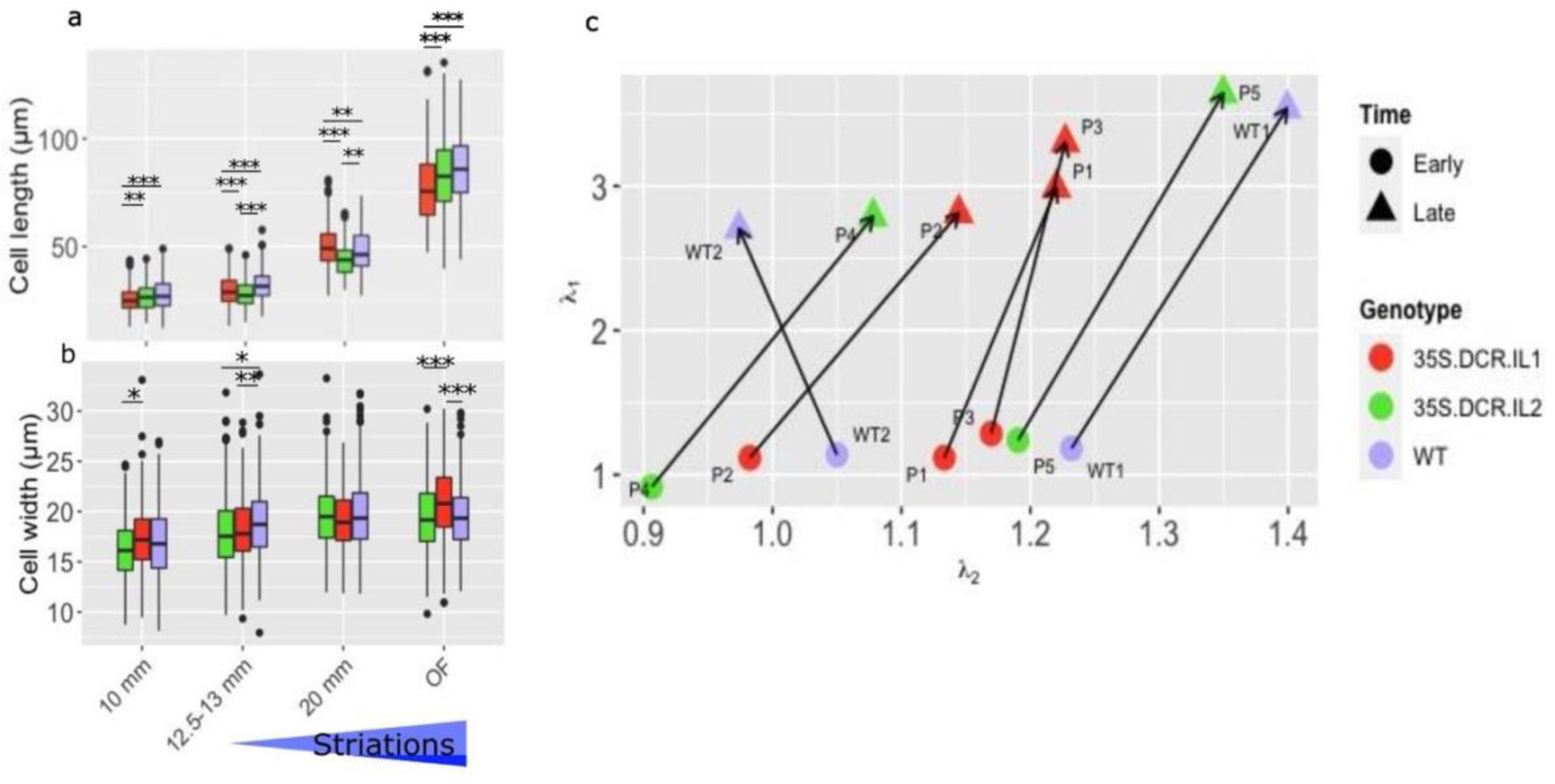
Overexpression of *HtDCR-like1* is not sufficient to alter the anisotropic growth of petal epidermal cells. **a** and **b**) Size of the adaxial proximal epidermal cells at different stages of development (10-, 12.5-13-and 20-mm buds as well as ‘open flower’ (OF)), measured in the length and width axes of the petals (**a** and **b**, respectively). Blue triangle represents the theoretical presence and development of striations in a WT individual (spanning from 12.5-13 mm buds to fully mature flowers). A Kruskal Wallis test was performed for each developmental stage and in case of p-value lower than 0.05 a subsequent pairwise Wilcoxon with Benjamini Hochberg’s correction test was done for a greater resolution. * ⇐⇒ 0.05 > p-value > 0.01, ** ⇐⇒ 0.01 > p-value > 0.001, *** ⇐⇒ 0.001 > p-value. **c**) Strains in the length (λ_1_) and width (λ_2_) axes of the petal calculated by dividing the cell size between 12.5-13- and 10-mm buds (initial stage(●)) and open flower and 10-mm buds (final stage(▴)).

Overall, wild type and transgenic samples encounter similar anisotropic cell deformation during the developmental stages at which buckling would normally occur. Hence, the failure of the buckling observed in the two transgenic lines is unlikely to be the result of an altered growth pattern of the epidermal cells.

### Overexpression of *HtDCR-like1* is sufficient to alter overall cuticle chemistry and results in lower detection of 10,16-DHP (C_16_H_32_O_4_)

The alteration of the cuticular striations in *HtDCR-like1* overexpression transgenic lines cannot be correlated with alteration of the cuticle thickness or with alteration of the growth of the epidermal cells. The potential impact of the overexpression of *HtDCR-like1* on the chemistry of the cuticle was therefore investigated using liquid extraction surface analysis coupled to high-resolution mass spectrometry (LESA-MS) (Giorio et al., 2015). The presence of 110 compounds was assessed in the adaxial cuticle film of WT *Hibiscus trionum* and *35S:HtDCR-like1* overexpression lines. The solvents used for liquid extraction do not allow for the hydrolysis of chemical bonds such as ester bonds and only enable the extraction of compounds that are free in the cuticular layer.

For each sample, a complex chemical profile was obtained. A principal component analysis was performed to highlight proximity between the samples based on their chemical profiles (Fig. **5a** and supplementary figure S4). Smooth distal petal parts and striated proximal petal parts clustered separately for all individuals, as found by (Moyroud et al., 2022). This suggests a relatively mild alteration of the chemistry of the adaxial proximal and distal petal cuticle in the transgenic lines. However, the distance between WT proximal and distal samples was considerably larger than that between the proximal and distal samples from the transgenic lines, which suggests that the overexpression of *HtDCR-like1* hinders the chemical differentiation of the proximal adaxial cuticle (see also supplementary figure S4). Since it has been suggested that AtDCR induces mid-chain hydroxylation of cutin monomers (Molina & Kosma, 2015), the detection rate of C_16_-C_18_ molecules in the cuticle was further investigated (**Fig. 5b** and **c** and supplementary figure S5). Only two compounds showed consistent differential detection between the WT and the transgenic lines of *Hibiscus trionum*: 10,16-dihydroxypalmitic acid (10,16-DHP/C_16_H_32_O_4_ see also supplementary table S2) and C_16_H_28_O_16_. An effect on detection of 10,16-DHP is consistent with previous research establishing the potential role of *AtDCR* in the regulation of 10,16-DHP (Panikashvili et al., 2009). Surprisingly, the detection frequency of 10,16-DHP was lower in *35S:HtDCR-like1* overexpression transgenic lines, suggesting that another molecular mechanism is involved in response to *HtDCR-like1* over expression and results in lower amounts of 10,16-DHP in the solubilised fraction of the cuticle. The second compound, C_16_H_28_O_16_, potentially corresponding to xylosylcellobiose, was detected more frequently in the transgenic lines than in the WT. Although its role and function remain unknown in the context of cuticle buckling, it has been shown in yeast that xylosylcellobiose is a by-product of xylosaccharide degradation by endoxylananases (Biely & Vršanská, 1983).

**Figure 5:**
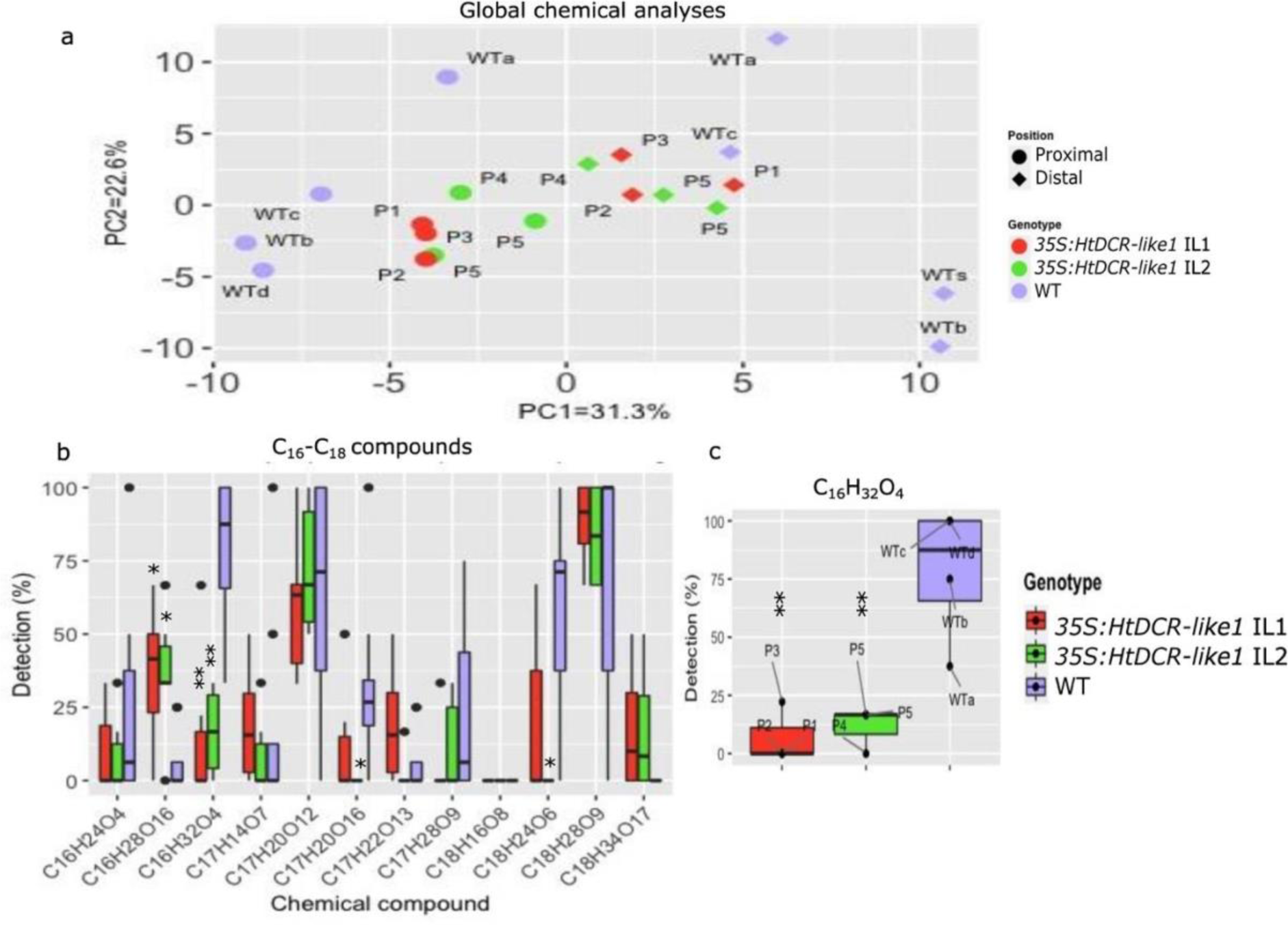
The *35S:HtDCR-like1* transgenic lines display an alteration of the chemistry of the adaxial cuticular film. **a**) Principal component analyses performed on the LESA-MS results accounting for the overall 110 compounds detected. **b**) Detection rate (i.e. proportion of replicate samples in which compound was later detected) observed for 17 chemical compounds of 16, 17 and 18 carbon residues performed on the proximal adaxial petal surface only. A Kruskal Wallis test was performed between the 3 samples for each compound. When p-value < 0.05, a subsequent pairwise Wilcoxon test was applied between the WT and each transgenic line with Benjamini and Hochberg’s correction. * ⇐⇒ 0.05 > p-value > 0.01, ** ⇐⇒ 0.01 > p-value > 0.001. **c**) Boxplot representing the percentage of C_16_H_32_O_4_ (10,16-DHP) detection for each genotype with the contribution of each sample. Four WT individuals were analysed using LESA-MS (samples a to d).

### Overexpression of *HtDCR-like1* is sufficient to alter the stiffness of the cuticle layers

Alteration of the cuticular composition, and especially of the cutin network, is likely to alter the mechanical properties of the cuticle (Domínguez et al., 2009). Furthermore, it has been suggested that the stiffness ratio of the proximal adaxial upper cuticle film and its substrate is critical to its buckling (Airoldi et al., 2021; Lugo et al., 2023). The Young’s modulus of the cuticle was measured during the initiation of buckling in WT petals (12.5-13 mm) and the equivalent developmental stage in the transgenic lines. Young’s moduli were assessed for the cuticle film (by shallow indentation on the intact tissue), and separately on the cuticular layer remaining after mechanical removal of the cuticle film. The Young’s modulus of the cuticle film could not be measured separately following its removal as the mechanical removal was destructive.

On average, the cuticular layer (the substrate) is stiffer in the two *35S:HtDCR-like1* transgenic lines than in the WT (Fig. **6**, Young’s moduli of 5.2 and 4.5 MPa for the transgenic lines and 2.3 MPa for the WT). Furthermore, in the WT petals, the stiffness of the cuticular layer is consistently below 7 MPa in all replicates, while it reaches values of over 15 MPa in the *HtDCR-like1* over expression lines. Meanwhile, the average stiffness measured on the upper cuticle film in the transgenic lines is lower than in the WT (23 and 16.4 MPa for the transgenic lines and 25.8 MPa for the WT). These data indicate that the transgenic lines have a stiffer substrate than the wild type and a softer upper layer. In combination, this results in a lower ratio of stiffness between the two (R, where R = Young’s modulus of upper layer/Young’s modulus of lower layer). Previous studies (Lugo et al., 2022) have shown that higher ratios of R are more likely to induce buckling in a bilayer system. We note that in the transgenic lines the stiffness of the two layers approaches similar values, which never occurs in the WT cuticle and would result in a failure of buckling according to model precisions (Lugo et al., 2022).

**Figure 6:**
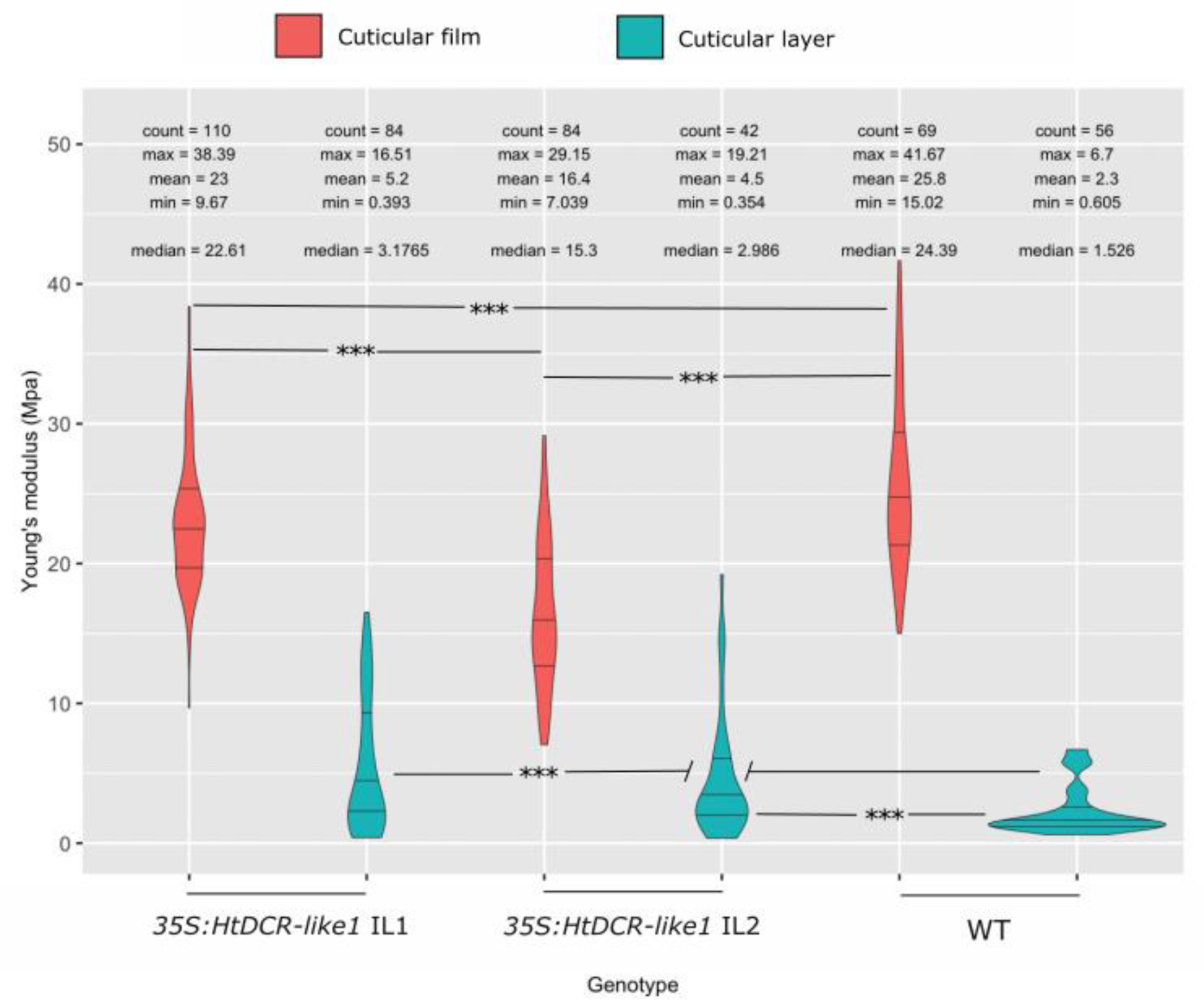
Over expression of *HtDCR-like1* is sufficient to alter cuticle stiffness. Violin plot of the Young’s moduli measured on the adaxial petal epidermis before and after peeling of the cuticular film in WT and *35S:HtDCR-like1* transgenic lines. Kruskal Wallis test showed significant differences in stiffness of both cuticle film and cuticular film and cuticular layer between the three genotypes. A pairwise Wilcoxon test was used subsequently for sample to sample comparison using Benjamini Hochberg’s correction: * ⇐⇒ 0.05 > p-value > 0.01, ** ⇐⇒ 0.01 > p-value > 0.001, *** ⇐⇒ 0.001 > p-value

The striations on mature transgenic flowers are patchily distributed and the AFM measurements were performed on buds prior to development of striations. The variation in our measurements is consistent with the observation that only parts of the cuticle exhibit the stiffness properties required for its buckling. It is likely that the cuticle film and the cuticular layer do not form two mechanically distinct layers with a sufficiently high stiffness ratio between them to allow cuticle buckling in all regions of the petals, but in some areas the mechanical properties are sufficient to allow buckling to occur.

## Discussion

In this article we demonstrate that overexpression of *HtDCR-like1* is sufficient to alter both the cuticle chemistry and the production of cuticular striations on the adaxial petal surface of *Hibiscus trionum.* We also show that the alteration of the cuticle chemistry results in modification of its Young’s modulus (stiffness). Consistent with the *Hibiscus trionum* cuticle buckling models (Airoldi et al., 2021 and Lugo et a., 2023), we show that reduction in the R-value (Young’s moduli ratio of the cuticle film (upper layer) and the cuticular lower layer) is sufficient to induce a failure of the cuticle buckling. Thus, we have linked together specific alteration of the cuticle chemistry with modulation of cuticle stiffness, confirming a potential way by which gene expression can control complex physical phenomena such as cuticular buckling.

Our finding also supports the potential key role of 10,16-dihydroxypalmitic acid in the development of cuticular striations as shown in *Arabidopsis thaliana* (Panikashvili et al., 2009), and as suggested in *Hibiscus trionum* (Moyroud et al., 2022). Surprisingly, despite the strong depletion of 10,16-DHP in the *dcr* mutants of *A. thaliana*, overexpression of *DCR* in *H. trionum* did not result in a higher detection of 10,16-DHP in *H. trionum,* but instead a depletion. It could be that an excess of 10,16-DHP was produced but was integrated in the cutin network and that it triggered higher reticulation levels, resulting in lower levels of loose 10,16-DHP. This hypothesis is consistent with a stiffer cuticular layer as shown in Fig. **6**.

The discovery that altering *HtDCR-like1* expression influences cuticular striations in *Hibiscus trionum* is not surprising, but the key link made here is to the stiffness of the cuticle. We measured partially overlapping Young’s moduli before and after mechanical removal of the cuticle in the two *35S:HtDCR-like1* transgenic lines, whereas in the WT, these values are markedly different. This suggests that in some instances, the R-value in the transgenic lines is close to 1. Mathematical and experimental models predict that an R-value of 1 leads to a failure of buckling (Airoldi et al., 2021; Cao & Hutchinson, 2012; Lugo et al., 2023). Therefore, it appears that alteration of the Young’s modulus is most likely the main reason for the failure of the buckling in the *35S:HtDCR-like1* transgenic lines.

Previous work has linked the accumulation of flavonoids in the cuticle to its elasticity during the ripening of tomato fruit (Domínguez et al., 2009). However, this previous study did not reveal any differential mechanical properties between the cuticle film and its substratum (or cuticular layer) upon the modification of its flavonoid content. Here, we demonstrate that overexpression of *HtDCR-like1* is sufficient to perturb the mechanical properties of the cuticle layers, reducing the difference in their Young’s moduli. Our finding also suggests that 10,16-DHP plays a central role in determining the mechanical properties of the cuticle and, thus, in the cuticle buckling and the production of structurally coloured cuticle. This hypothesis is supported by the differential amounts of 10,16-DHP that were measured between striated and non-striated cuticles in various *Hibiscus* species and varieties and transgenic lines devoid of striations (Moyroud et al., 2022), and by the depletion of 10,16-DHP that was observed in the *defective in cuticular ridges* (*dcr*) mutant of *A. thaliana* (Panikashvili et al., 2009).

The alteration of cell wall compounds has not yet been investigated as a means to achieve buckling in the epidermal cuticle. However, cellulose and pectin often populate the deepest layers of the epidermal cuticular layer. It can be speculated that the Young’s modulus of these deeper layers could be adjusted by modulation of the reticulation of the cellulose network, or by modulation of the pectin properties. This could represent an essential parameter in the softening of the substratum below the cuticular film and may constitute another route to achieving cuticule buckling.

The production of iridescent cuticles is specific to epidermal tissues composed of flat cells. On the adaxial petal cuticle of *Hibiscus trionum*, the occurrence of flat elongated cells is correlated with the presence of nano-scaled cuticular ridges. The distal adaxial petal region is characterised by both a change in cell shape and a loss of striations. It can thus be speculated that cell shape and cuticle chemistry are tightly co-regulated to enable the production of structurally coloured surfaces. Transcriptomic analyses may help us to further understand the early transcription factors and effectors involved in the regulation of cell type and cuticle patterning.

In this study we elucidate one of the mechanisms by which the production of structurally coloured cuticle is achieved on the petal of *Hibiscus trionum* by linking together the regulation of *HtDCR-like1*, the chemistry of the cuticle, the mechanical properties of the cuticle and its capacity to buckle. We show that the regulation of *HtDCR-like1* controls the chemistry of the cuticle and plays a specific role in determining the cuticle content in 10,16-dihydroxypalmitic acid. We demonstrate that specific alteration of the cuticle chemistry leads to an alteration of the Young’s moduli of the cuticle layers and propose that such a modification is sufficient to impair the buckling of the cuticle, resulting in partial loss of the structural colour of *Hibiscus trionum* flowers.

## Acknowledgments

This work was funded by BBSRC grants BB/P001157/1 and BB/V000314/1 to BJG and by the PlaMatSu identifier H2020-MSCA-ITN-2016 No 722842. CryoSEM images and AFM measurements were acquired thanks to the cryoSEM and AFM platforms at the Microscopy Core Facility of the Sainsbury Laboratory Cambridge University which is supported by the Gatsby Charitable Foundation. Liquid extraction surface analysis coupled with high-resolution mass spectrometry experiments were supported by a BP Next Generation fellowship awarded by the Yusuf Hamied Department of Chemistry at the University of Cambridge to Chiara Giorio. Specific acknowledgements go to Matthew Dorling for excellent plant care, Qi Wang for bioinformatic support and Carlos Lugo-Velez and Edwige Moyroud for stimulating discussions leading to the production of this paper.

## Authors’ contribution

Jordan Ferria: experimental design, paper writing, generation of transgenic lines, phylogeny work, chemical data analyses, cell size data acquisition and analysis, Atomic Force Microscopy data analysis, Cryo-SEM image acquisition and analysis. Siriel Saladin: liquid extraction surface analysis coupled with high-resolution mass spectrometry experiments, data analysis and paper reviewing. Udhaya Ponraj: Atomic Force Microscopy data acquisition and paper reviewing. Raymond Wightman: Cryo-SEM sample preparation and image acquisition. Chiara Giorio: supervision of liquid extraction surface analysis coupled with high-resolution mass spectrometry experiments and paper reviewing. Chiara Airoldi: experimental design, supervision and paper reviewing. Beverley Glover: experimental design, supervision, paper reviewing and funding acquisition.

---

**Supplementary table S1:**
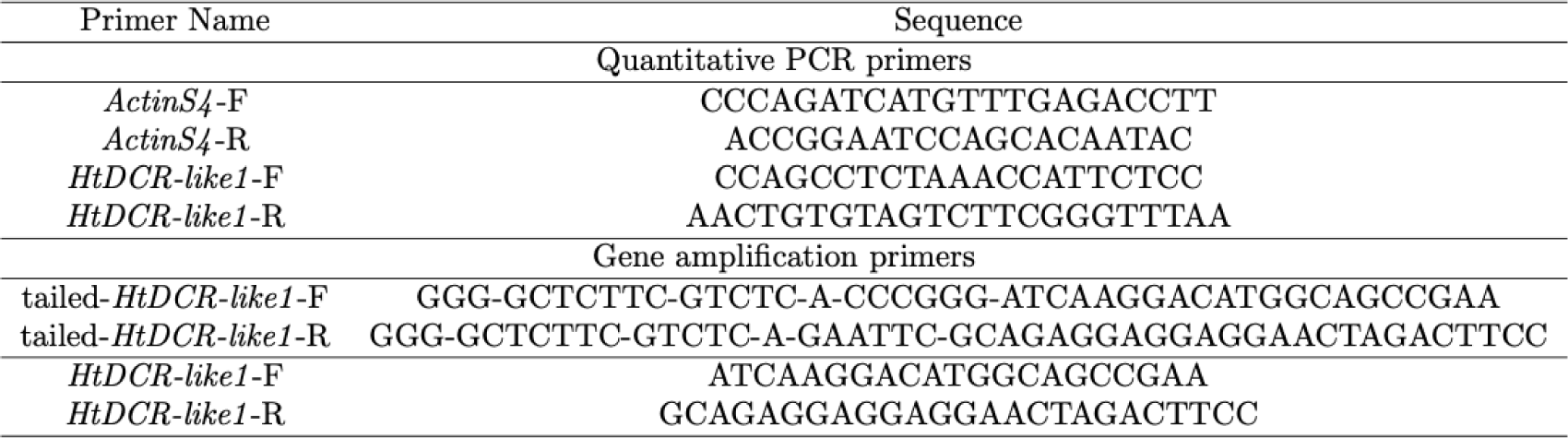
Sequence of the primers used for the amplification of *HtDCR-like1* from cDNA and for quantitative PCR in the WT and *35S:HtDCR-like1* transgenic lines.

**Supplementary table S2:**
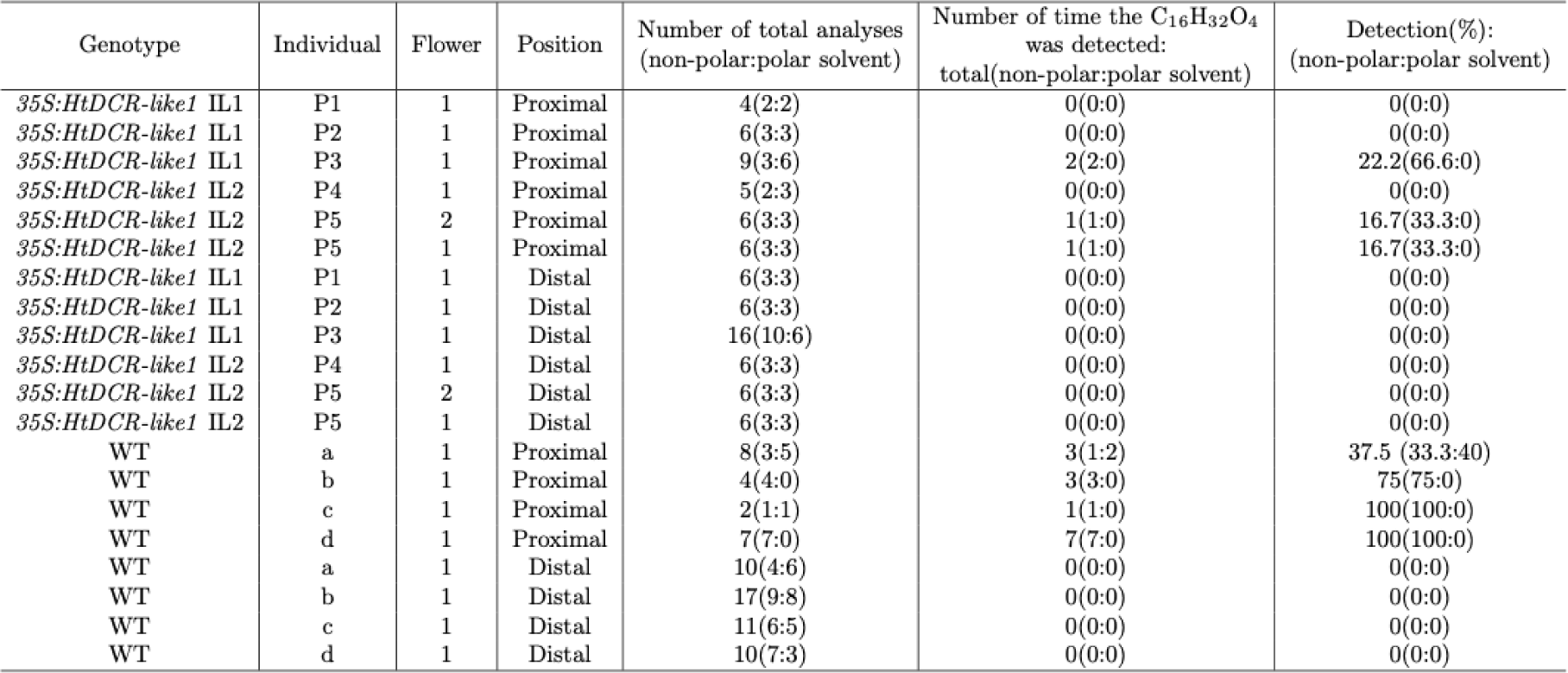
Chemical data collected from WT and *35S:HtDCR-like1* transgenic lines filtered for the C_16_H_32_O_4_ compound.

| Genotype | Individual | Flower | Position | Number of total analyses<br>(non-polar:polar solvent) | Number of time the C<sub>16</sub>H<sub>32</sub>O<sub>4</sub><br>was detected:<br>total(non-polar:polar solvent) | Detection(%):<br>(non-polar:polar solvent) |
| --- | --- | --- | --- | --- | --- | --- |
| *35S:HtDCR-like1* IL1 | P1 | 1 | Proximal | 4(2:2) | 0(0:0) | 0(0:0) |
| *35S:HtDCR-like1* IL1 | P2 | 1 | Proximal | 6(3:3) | 0(0:0) | 0(0:0) |
| *35S:HtDCR-like1* IL1 | P3 | 1 | Proximal | 9(3:6) | 2(2:0) | 22.2(66.6:0) |
| *35S:HtDCR-like1* IL2 | P4 | 1 | Proximal | 5(2:3) | 0(0:0) | 0(0:0) |
| *35S:HtDCR-like1* IL2 | P5 | 2 | Proximal | 6(3:3) | 1(1:0) | 16.7(33.3:0) |
| *35S:HtDCR-like1* IL2 | P5 | 1 | Proximal | 6(3:3) | 1(1:0) | 16.7(33.3:0) |
| *35S:HtDCR-like1* IL1 | P1 | 1 | Distal | 6(3:3) | 0(0:0) | 0(0:0) |
| *35S:HtDCR-like1* IL1 | P2 | 1 | Distal | 6(3:3) | 0(0:0) | 0(0:0) |
| *35S:HtDCR-like1* IL1 | P3 | 1 | Distal | 16(10:6) | 0(0:0) | 0(0:0) |
| *35S:HtDCR-like1* IL2 | P4 | 1 | Distal | 6(3:3) | 0(0:0) | 0(0:0) |
| *35S:HtDCR-like1* IL2 | P5 | 2 | Distal | 6(3:3) | 0(0:0) | 0(0:0) |
| *35S:HtDCR-like1* IL2 | P5 | 1 | Distal | 6(3:3) | 0(0:0) | 0(0:0) |
| WT | a | 1 | Proximal | 8(3:5) | 3(1:2) | 37.5 (33.3:40) |
| WT | b | 1 | Proximal | 4(4:0) | 3(3:0) | 75(75:0) |
| WT | c | 1 | Proximal | 2(1:1) | 1(1:0) | 100(100:0) |
| WT | d | 1 | Proximal | 7(7:0) | 7(7:0) | 100(100:0) |
| WT | a | 1 | Distal | 10(4:6) | 0(0:0) | 0(0:0) |
| WT | b | 1 | Distal | 17(9:8) | 0(0:0) | 0(0:0) |
| WT | c | 1 | Distal | 11(6:5) | 0(0:0) | 0(0:0) |
| WT | d | 1 | Distal | 10(7:3) | 0(0:0) | 0(0:0) |

**Supplementary figure S1:**
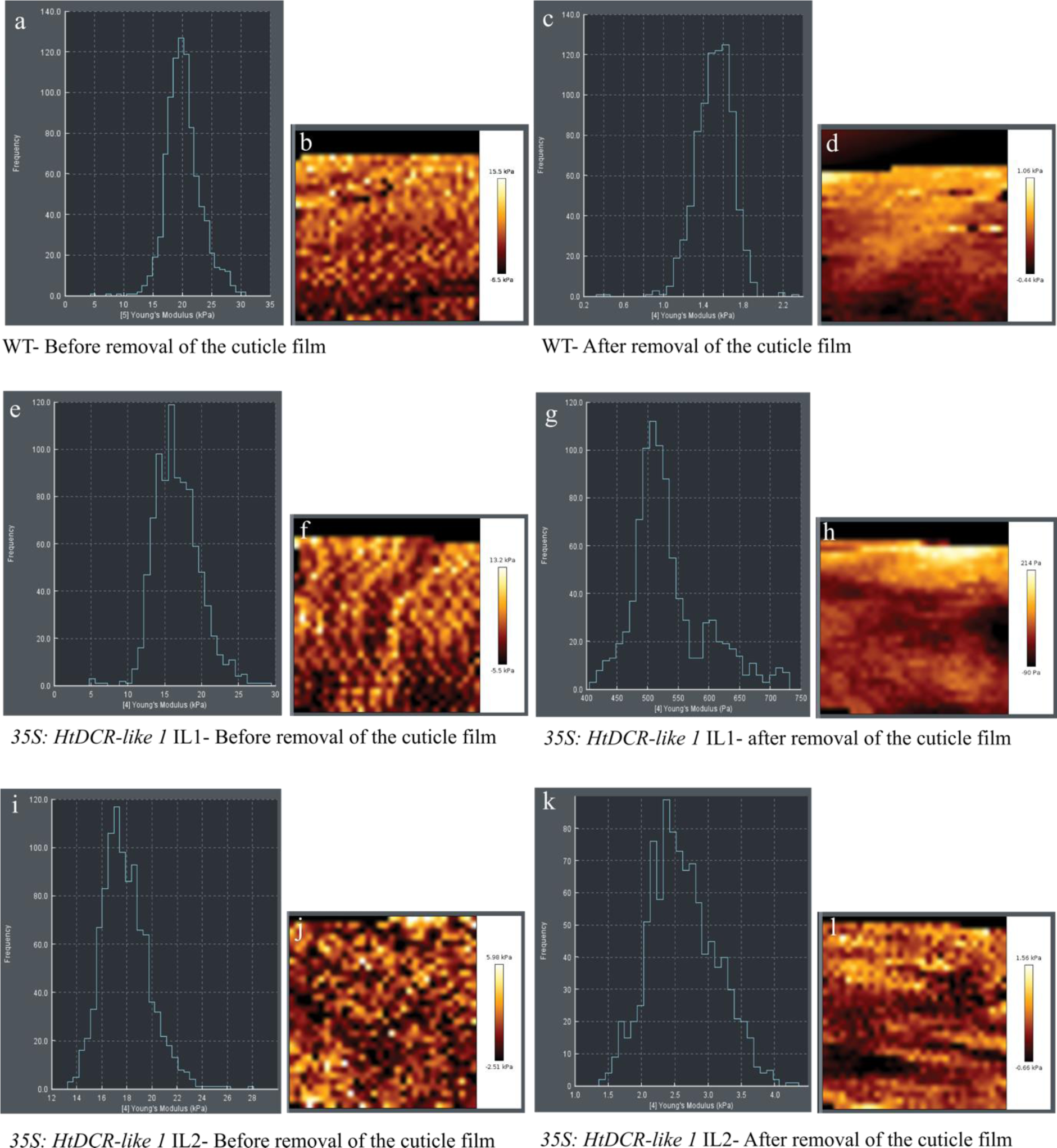
Measurement of the cuticle stiffness using Atomic Force Microscopy (AFM). **a**, **c**, **e**, **g**, **i** and **k**) Examples of frequency curves obtained from force displacement curves measured with a grid of 32 x 32 points for one sample of WT, *35S:HtDCR-like1* IL1 and *35S:HtDCR-like1* IL2 transgenic lines. **b**, **d**, **f**, **h**, **j** and **l**) 20 μm x 20 μm AFM maps of Young’s moduli used to produce the frequency curves. **a**, **b**, **c** and **d**) WT. **e**, **f**, **g** and **h**) *35S:HtDCR-like1* Independent line 1. **i**, **j**, **k** and **l**) *35S:HtDCR-like1* Independent line 2. **a**, **b**, **e**, **f**, **i** and **j**) Before mechanical removal of the cuticle film. **c**, **d**, **g**, **h**, **k** and **l**) After mechanical removal of the cuticle film.

**Supplementary figure S2:**
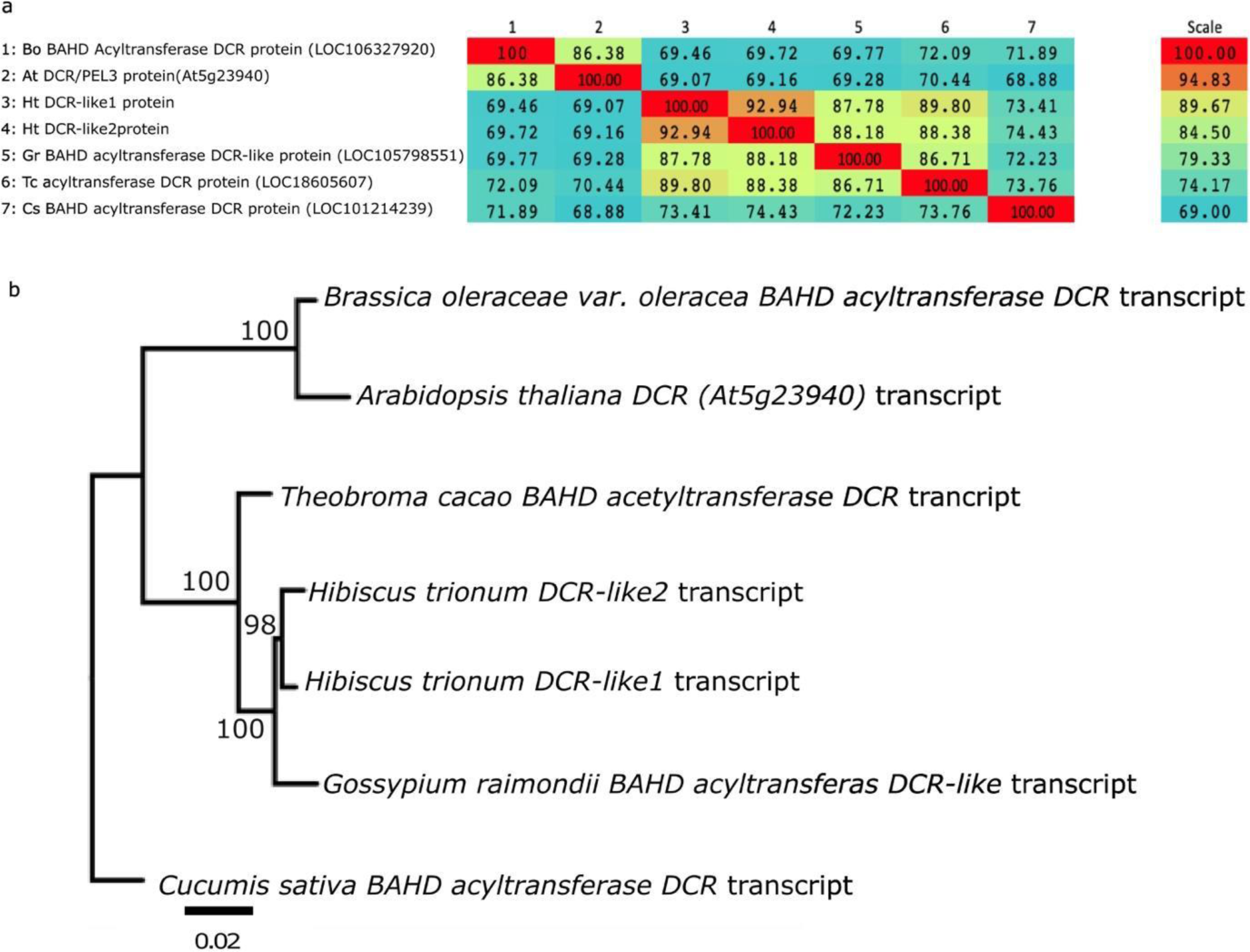
**a**) Percentage of identity between protein sequences based on the protein sequence alignment displayed in figure 1. **b**) Bootstrapped phylogeny based on *DCR* and *DCR-like* RNA sequences using the JTT model of raxml (similar results were obtained using the DCMUT+G model, data not shown)

**Supplementary figure S3:**
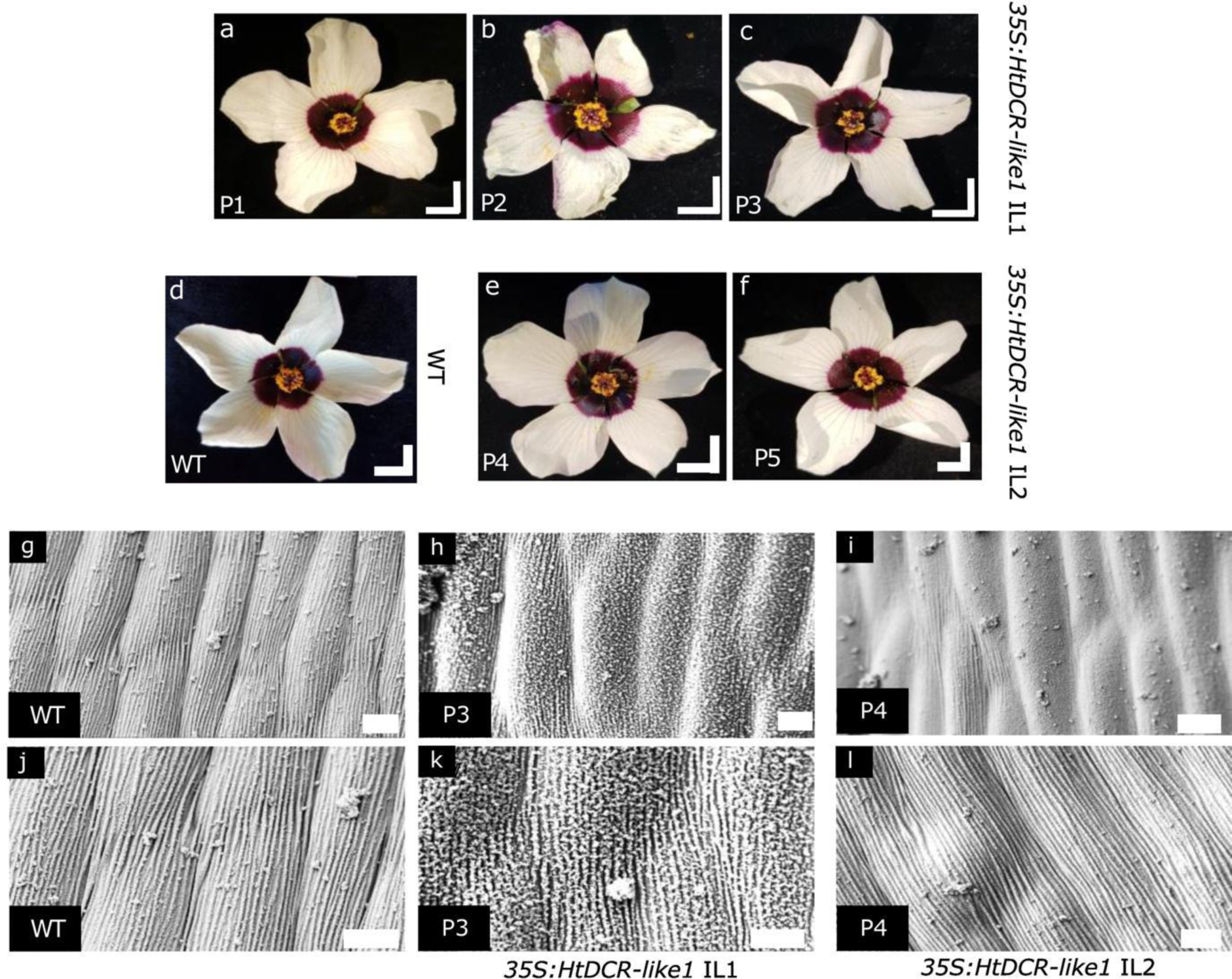
Macroscopic pictures and scanning electron microscopy of WT and *35S:HtDCR-like1* transgenic lines. **a-f**) Flower macroscopic pictures (scale bar = 1 cm). **g-l**) SEM of adaxial proximal petal surface (scale bar = 10μm). **a**, **b**, **c**, **h** and **k**) 35S:HtDCR-like1 IL1transgenic line. **e**, **f**, **i** and **l**) *35S:HtDCR-like1* IL2 transgenic line. **d**, **g**, **j**) WT. **a**) P1 plant, b P2 plant, **c**, **h**, **k**) P3 plant. **e**, **i**, **l**) P4 plant, **f**) P5 plant.

**Supplementary figure S4:**
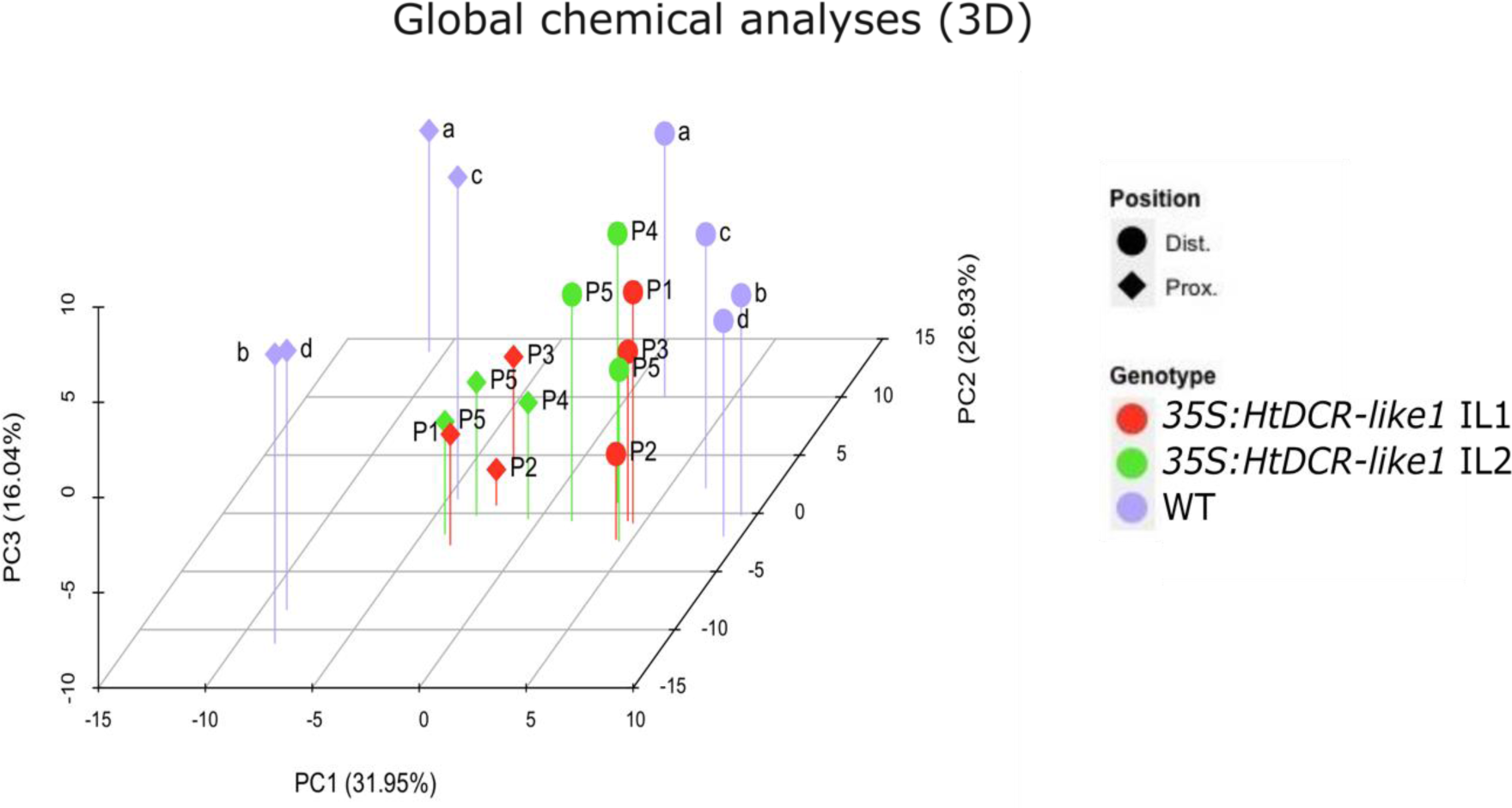
3D-Principal component analysis performed on the LESA-MS results accounting for the overall 110 compounds detected ccounting for about 67% of the total variance.

**Supplementary figure S5:**
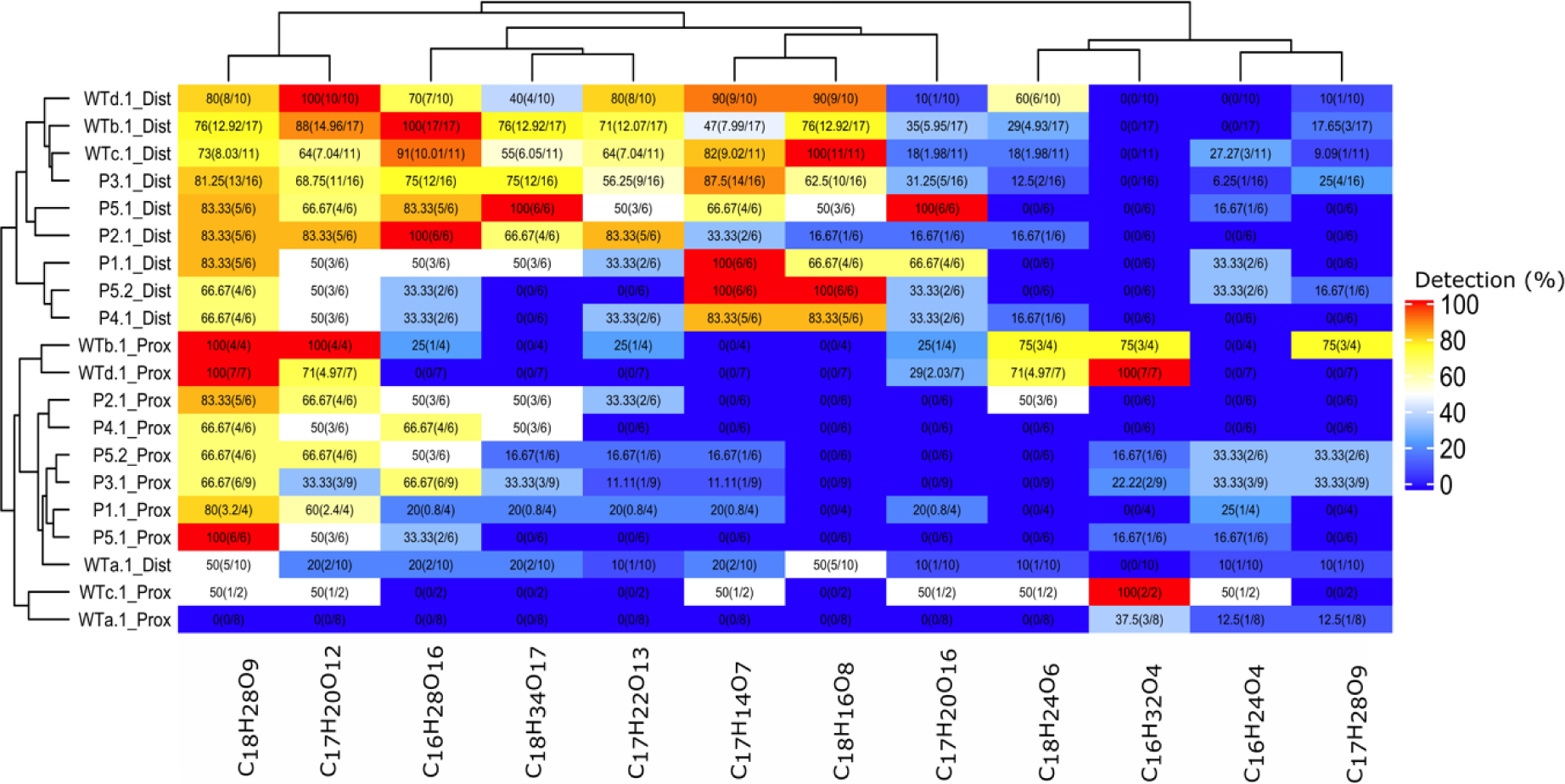
Heatmap summarising the detection rate measured for each of the chemical compounds composed of 16, 17 and 18 carbon residues as detected by LESA-MS, expressed in percentage (before the brackets). c (a/b) ⇐⇒ a: the number of times the compound was detected. b: the number of detection attempts. c: the ratio of ‘a’ and ‘b’ multiplied by 100. ‘Distal’ refers to the white, distal and non-striated region of the adaxial petal part. ‘Proximal’ refers to the proximal striated (at least in the WT samples) region of the adaxial petal part. P(1-5) and letters from a to d refer to individual plants. Numbers 1 and 2 refer to the flower that was analysed

## Notes

### Competing Interest Statement

The authors have declared no competing interest.

